# Punishing temporal judgement boosts sense of agency and modulates its underlying neural correlates

**DOI:** 10.1101/2024.12.12.628000

**Authors:** Christopher M. Hill, Numa Samnani, Leo Barzi, Matt Wilson

## Abstract

Feeling in control of one’s actions is fundamental to the formation of action-outcome relationships. Reinforcement and its valence also change the action-outcome relationship, either through behavior promotion or diminishment. In this study we evaluated how reward and punishment reinforcement modulate sense of agency, as measured by intentional binding. Moreover, using electroencephalography (EEG) we evaluated how reward and punishment reinforcement changes outcome event related potentials associated with the accuracy of participants’ judgement of the time interval between a key press and audio tone. We found that punishment reinforcement increased intentional binding between the action and outcome more than reward and control feedback. This was also reflected in the outcome event related potentials, where punishment elicited greater P300s and Late Positive Potentials compared to reward and control. We also found increased N100s and diminished P300s and Late Positive Potentials when the participants did not actively participate in evoking the tone. Taken together, our findings showcase that punishment reinforcement boosts sense of agency and modulates associated neural activity more than reward and no reinforcement, as a function of increasing attention and arousal. These findings illuminate the greater effect punishment reinforcement has on behavior and brain activity by its modification of sense of agency, which is important for the development of treatments in psychiatric and neurological diseases.

## Introduction

When we perform voluntary actions, there is a sensation of ownership that corresponds with their execution, known as the “sense of agency”. This subjective experience has been widely studied in cognitive science especially in the context of consciousness, and free will [1,2]. Changes in sense of agency have been noted in many psychiatric and neurological disorders [1–4]. One widely used measure of sense of agency is the intentional binding effect, the temporal perception of two events [4–6], where voluntary actions and their consequences are perceived as being closer together and involuntary actions and their consequences are perceived more independently. Often intentional binding is assessed by comparing a person’s temporal perception between self-initiated and passive actions [1,5,7], where a person judges how much time passed between an action (i.e. button press) and an audio tone [4–6].

Previous behavioral investigations have examined the relationship between outcome valence and sense of agency. Modulating outcome valence is typically done by making the tone heard after the action, more appetitive or aversive, either by changing the tone frequency or pairing it with a visual stimulus. However, there have been mixed results regarding outcome valence and the intentional binding effect. For instance, studies show aversive outcome tones increase intentional binding [8,9], while others found decreases [10,11]. Appetitive tones also decreases the intentional binding effect [12], but may be more effective than aversive tones by comparison [10,11]. Moreover, some studies have found no differences in between appetitive and aversive outcomes [13,14]. Given the mixed nature of these findings, the effect of outcome valence on intentional binding is not clearly understood and needs further exploration [15].

Reinforcement feedback is a powerful tool in shaping the relationship between our actions and their associated outcomes to promote behavioral change [16–19]. Two common forms of reinforcement feedback are reward and punishment, which promote or diminish occurrences of certain behaviors, in turn strengthening action-outcome relationships [16–19]. However, reinforcement feedback in the context of sense of agency has not been extensively studied and has largely remained separate [20]. One recent investigation evaluated how reinforcement feedback based on action timing accuracy, found that intentional binding decreased when accuracy increased, as a function of task learning and lowering cognitive load during action selection [20].

Electroencephalography (EEG) has been used to evaluate brain activity during temporal perception tasks. Specifically, outcome event-related potentials (ERPs) have provided valuable insights into the cortical processing related to sense of agency. Distinct components of outcome ERPs peak between 50 and 600 ms after task relevant stimuli and reflect different aspects of outcome monitoring, including attention, stimuli significance, and saliency [21–25]. Previous studies demonstrate early components of outcome ERPs decrease in amplitude when a person voluntarily generates an outcome versus externally generated [26–28]. Instances of coercion and manipulation (i.e. lower agency) decreases the amplitude of early outcome ERP components in comparison to free choice [29,30]. Conversely, amplitudes of later outcome ERP components increase in conditions where participants have higher control over their actions compared to low or no control [26,27,31].

Outcome ERPs have also been studied with reinforcement feedback across a wide variety of contexts [16,22,32–40]. Reward and punishment demonstrate differential effects on outcome ERPs thus reflecting different brain processing. Punishment elicits greater amplitude in early outcome ERP components [41]. In part, early components represent activity in the anterior cingulate cortex, a portion of prefrontal cortex responsible for the detection of error and performance monitoring [16,32,33,37,42]. Later outcome ERP components have demonstrated increases with both reward and punishment feedback, reflecting greater attention and saliency [41,43,44].

Studies evaluating sense of agency via intentional binding have primarily focused on the outcome valence (i.e. tone) after a voluntary or involuntary action [9,12,20]. A critical component of determining intentional binding is the temporal judgement made by the participant, where they report how much time they thought passed between the action and outcome [1,4,45]. However, there is a lack of studies evaluating how feedback related to temporal judgements can modulate intentional binding and its associated neural activity. As a result, there is an open question about how the feedback valence in response to our temporal judgements uniquely changes our sense of agency and underlying brain processes. To this end, we utilized a temporal estimation task to examine how reward and punishment feedback based on temporal judgement accuracy changes the intentional binding effect and outcome ERPs in response to reinforcement feedback provided after their temporal judgements.

## Methodology

All procedures were approved by the University Institutional Review Board and were conducted according to the principles expressed in the Declaration of Helsinki. All participants provided written informed consent prior to participating. Thirty-three young healthy adults participated in this study [age range: 19–30 years, mean age ± standard deviation (SD): 21.84 ± 2.52 years, Edinburgh Handedness Inventory (EHI) mean handedness score ± SD: 66.90 ± 55.99, males: 18, females: 15)]. Participants were free of major physiological (musculoskeletal, neurological, cardiovascular) and psychological (drug abuse, depression) disorders. Participants were recruited from the local population of the University and the surrounding communities using word of mouth, electronic announcements, and posted flyers. Each participant was randomly allocated to one of three feedback groups [Reward (n=11, males=8, females=3), Punishment (n=11, males=3, females=8), Control (n=11, males=7, females=4)]. The Behavioral Avoidance/Inhibition scales (BAS/BIS) were used to score sensitivity to reinforcement which is divided into four subcomponents (BAS FUN, BAS DRIVE, BAS REWARD RESPONSIVENESS, BIS). Additional information regarding these scales is available elsewhere [46].

### Experimental procedures

#### Interval estimation paradigm

Participants were seated in a chair in front of a computer monitor, where they engaged in a modified version of Siebertz and Jansen (2022)’s interval estimation-paradigm via PsychoPy® software. The interval estimation-paradigm consisted of three blocks: one interval estimation-practice block (twelve practice trials total), followed by two counter-balanced active interval (90 active trials total) and passive interval (90 passive trials total) estimation blocks. During each practice trial, a fixation cross and the word ‘ready’ were displayed to prompt the participant to press the spacebar and trigger two sinusoidal tones (880 Hz), lasting 50 milliseconds (ms) each. The tones were presented binaurally via insert earphones (Etymotic Research, Elk Grove Village, IL, model ER-3A). The first tone was presented 200ms after the key press and a second tone was presented at one of three base intervals (200, 400, or 800ms) with a modified random jitter ±100ms. Participants used the number row on a keyboard to enter their estimation of the interval between the tones and pressed enter. The actual interval in milliseconds was then displayed, followed by neutral feedback displaying the difference between the participant input and actual interval. Practice trials were not analyzed [6].

After the practice block, participants engaged in active and passive interval estimation tasks, counter-balanced for each participant. In these blocks pressing the spacebar triggered only one tone, and participants were asked to estimate the interval between the key press and the tone which was presented at one of three base intervals (200, 400, or 800ms) with a modified random jitter ±100ms. 500ms after the tone, the prompt to enter their estimation was presented without feedback given (Fig 1). Each base interval was randomly presented for 30 trials (90 trials total per condition). In the active condition, participants pressed the button themselves, and in the passive condition, participants observed the experimenter pressing the button. The experimenter sat in a chair beside the participants in both conditions but did not engage during the active condition [6]. This same experimenter performed all experiments. For both conditions, a participant’s mean judgement error was calculated by subtracting the actual from the estimated intervals and presented on screen. A positive judgement error indicated overestimation, and a negative judgement error indicated underestimation of the temporal intervals [6].

**Fig 1:**
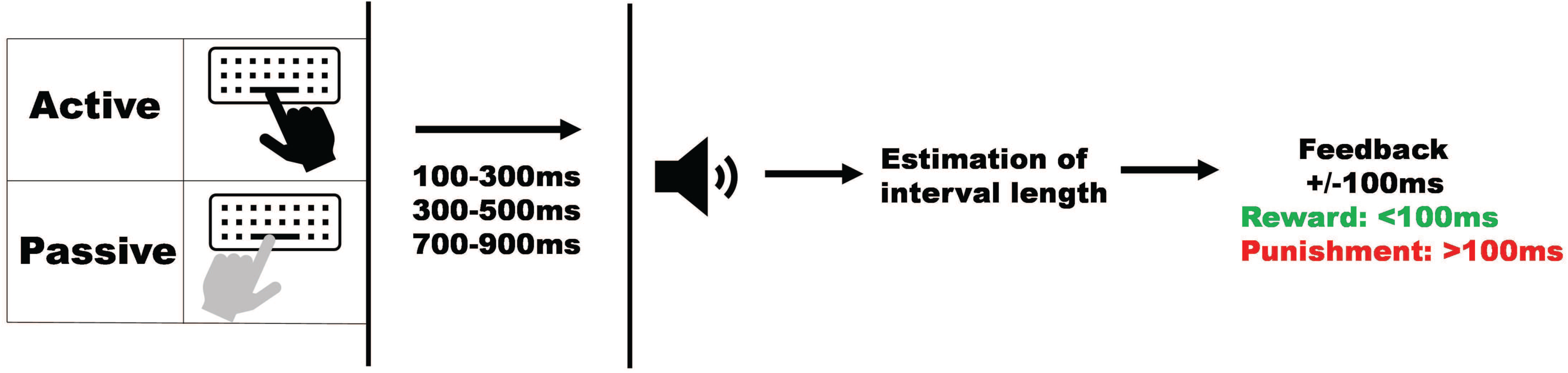
A schematic of the experimental procedures. The participant (Active) or the experimenter (Passive) pressed the button. This generated a tone after one of the three base time intervals. Participants then input their time estimation between the button press and the tone. Feedback was provided based on the difference between participants estimation and the actual tone latency.

Each person was allocated to one of three groups: Reward, Punishment, and Control. The color of the presentation of the judgement error determined the group assignment. The Reward feedback group was presented with green judgement error when the judgement error was within 100ms of the actual interval (Target Feedback) and white judgement error when not within 100ms of the actual interval (Neutral Feedback). The Punishment feedback group was presented with red judgement error when the judgement error was not within 100 ms of the actual interval (Target Feedback) and white judgement error when within 100ms of the actual interval (Neutral Feedback). The Control feedback group was presented with white judgement error regardless of the magnitude of the difference between the estimated and actual intervals.

Grand average outcome ERP waveforms were computed for each participant in each action condition (Active, Passive) and feedback type (Target, Neutral). Outcome ERP waveforms were visually inspected to identify waveform components [50] corresponding with previous intentional binding [28,29,31] and reinforcement feedback [33,34,51–58] studies. We identified waveform components N100, P300, and Late Positive Potential and analyzed peak amplitude and latency for all groups and conditions by looking in the following latency ranges: N100 (50 – 170 msec) [26,27], P300 (300 – 500 msec) [51–53], Late Positive Potential (515 – 650 msec) [56,58,59]. Amplitude and latency analyses were performed in electrodes where specific waveform components are most prominent. N100 amplitude and latency was extracted from the Fz electrode [32,60]. Whereas, P300 and Late Positive Potential amplitudes and latencies were obtained from the Pz electrode [51–53,58,61].

### Statistical analysis

Participant descriptive data and BAS/BIS subscales were evaluated with separate One-way analysis of variances (ANOVAs) to determine differences between the three feedback groups (Control, Reward, Punishment).

Mixed repeated measures ANOVAs were used to determine differences in judgement error and intentional binding across Feedback Groups (Reward, Punishment, Control) and Time Intervals (200, 400, 800). Violations of sphericity were corrected using the Greenhouse-Geisser correction factor.

Separate linear mixed models for repeated measures were used to determine differences for ERP amplitudes and latencies across feedback groups, feedback type, and action conditions. ERP amplitudes and latencies were held as the dependent variables. Feedback Group (Reward, Punishment, Control), Feedback Type (Target, Neutral), and Action Condition (Active, Passive) were held as fixed effects and individual subjects were held as random effects [62–65]. All follow-up analysis for main effects and interactions were performed with a Sidak correction. All statistical analysis was conducted in SPSS v26 (IBM Corp., Armonk, NY, United States).

## Results

### Participant characteristics

All groups demonstrated similar descriptive characteristics (Table 1). A One-way ANOVA revealed there was no significant differences among the three testing groups for EHI (F(2,30)=0.392, p=0.679) and Age (F(2,30)=0.587, p=0.562). All groups demonstrated similar sensitivities to reinforcement feedback. A One-way ANOVA revealed there was no significant difference among the three testing groups on the four subscales within the BAS/BIS scale [BAS Drive (F(2,30)=0.070, p=0.932), BAS FUN (F(2,30)=0.599, p=0.556), BAS Reward Responsiveness (F(2,30)=0.471, p=0.629), BIS (F(2,30)=0.260, p=0.772)].

**Table 1.** Participant Descriptive Data. Presented as mean ± standard deviation.

| Group | Age (Years) | EHI | BAS DRIVE | BAS FUN | BAS REWARD | BIS |
| --- | --- | --- | --- | --- | --- | --- |
| REWARD | 21.27 ± 0.13 | 54.45 ± 6.18 | 11.00 ± 0.19 | 11.63 ± 0.22 | 17.72 ± 0.14 | 21.09 ± 0.35 |
| PUNISHMENT | 22.45 ± 0.33 | 73.23 ± 4.94 | 11.18 ± 0.18 | 12.00 ± 0.15 | 18.45 ± 0.19 | 22.18 ± 0.19 |
| CONTROL | 21.81 ± 0.26 | 73.00 ± 5.92 | 10.81 ± 0.28 | 12.55 ± 0.21 | 18.18 ± 0.18 | 21.18 ± 0.41 |

### Judgement error

Judgement error increased as the time interval increased in both the Active and Passive conditions (Fig 2A). A significant main effect was found for Interval (F(2,60)=18.921, p<0.001) in the Active condition. Post-hoc comparisons revealed lower judgement error in interval 200 compared 400 [MD: 28.013, p=0.015, 95% CIs= 0.471-55.555] and 800 [MD: 68.417, p<0.001, 95% CIs= 40.875-95.959]. Moreover, a lower judgement error was found in interval 400 compared to 800 [MD: 40.404, p=0.002, 95% CIs= 12.862-67.946]. A significant main effect for Interval (F(2,60)=18.321, p<0.001) for the Passive condition. Post-hoc comparisons revealed lower judgement error in interval 200 compared 400 [MD: 56.957, p=0.001, 95% CIs= 28.891-85.022] and 800 [MD: 62.171, p<0.001, 95% CIs= 34.106-90.236]. No significant differences were found between intervals 400 and 800 [MD: 5.214, p=0.649, 95% CIs= −33.279-22.851].

**Fig 2:**
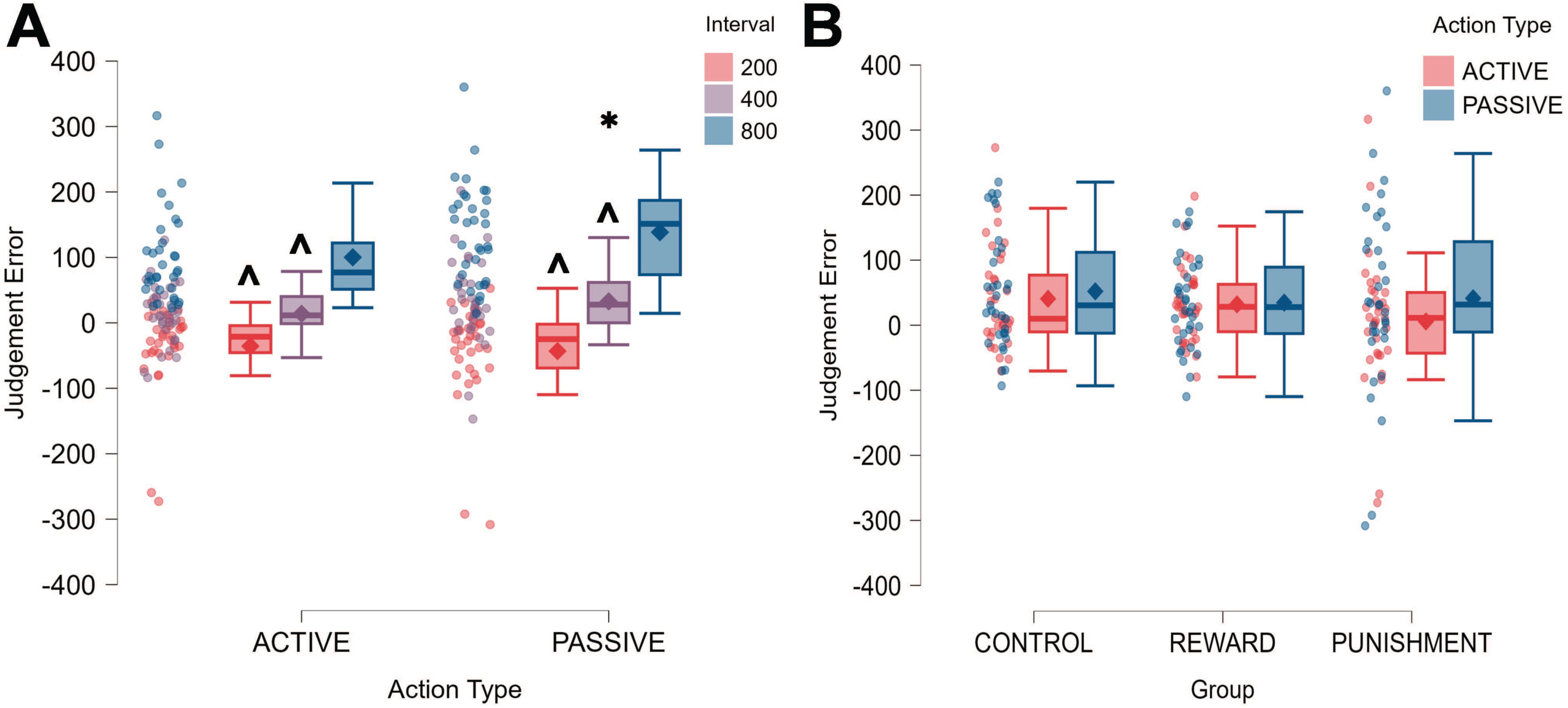
**A.** Boxplots for Judgement Error for each time interval [200ms (red), 400ms (purple), 800ms (blue)] in the Active and Passive conditions. Box ends represent the interquartile range (IQR) [25^th^ percentile (Q1) −75^th^ percentile (Q3)]. Whiskers represent minimum (Q1-1.5*IQR) and maximum (Q3 – 1.5*IQR). Median is represented by horizontal line inside of the box. Diamonds represent means and dots represent individual data. * = significantly different from Active. ^ = significantly different from 800ms. **B**. Boxplots for Reward, Punishment, and Control groups’ Judgement Error in the Active (red) and Passive (blue) conditions. Diamonds represent means and dots represent individual data

Average judgement error was different when the participant pressed the space bar than when the experimenter pressed it. A significant main effect for Condition (F(1,30)=8.988, p=0.005) was found where average judgement error during the Passive condition was higher compared to the Active condition [MD: 18.213, p=0.005, 95% CIs= 5.806-30.620]. No significant differences were found for between Groups (p>0.05) (Fig 2B).

### Intentional binding

Intentional binding was dependent on the time interval and the assigned group. A significant main effect for Interval (F(1.229,36.863)=21.550, p<0.001) was found (Fig 3). Post-hoc comparisons revealed lower intentional binding in interval 800 compared 200 [MD: 61.860, p=0.005, 95% CIs= 0.084-1.702] and 400 [MD: 100.172, p<0.001, 95% CIs= 0.789-2.104]. Similarly, there was lower intentional binding in interval 200 compared to 400 [MD: 38.312, p=0.007, 95% CIs= 0.095-1.011]. A significant main effect for Feedback Type (F(2,30)=3.526, p=0.042) was found. Punishment elicited greater intentional binding across all intervals compared to Reward [MD: 52.588, p=0.039, 95% CIs= 2.017-103.159] but not Control [MD: 31.747, p=0.244, 95% CIs= 18.823-82.318]. No differences were found between Control and Reward [MD: 20.840, p=0.304, 95% CIs= −29.730-71.411].

**Fig 3:**
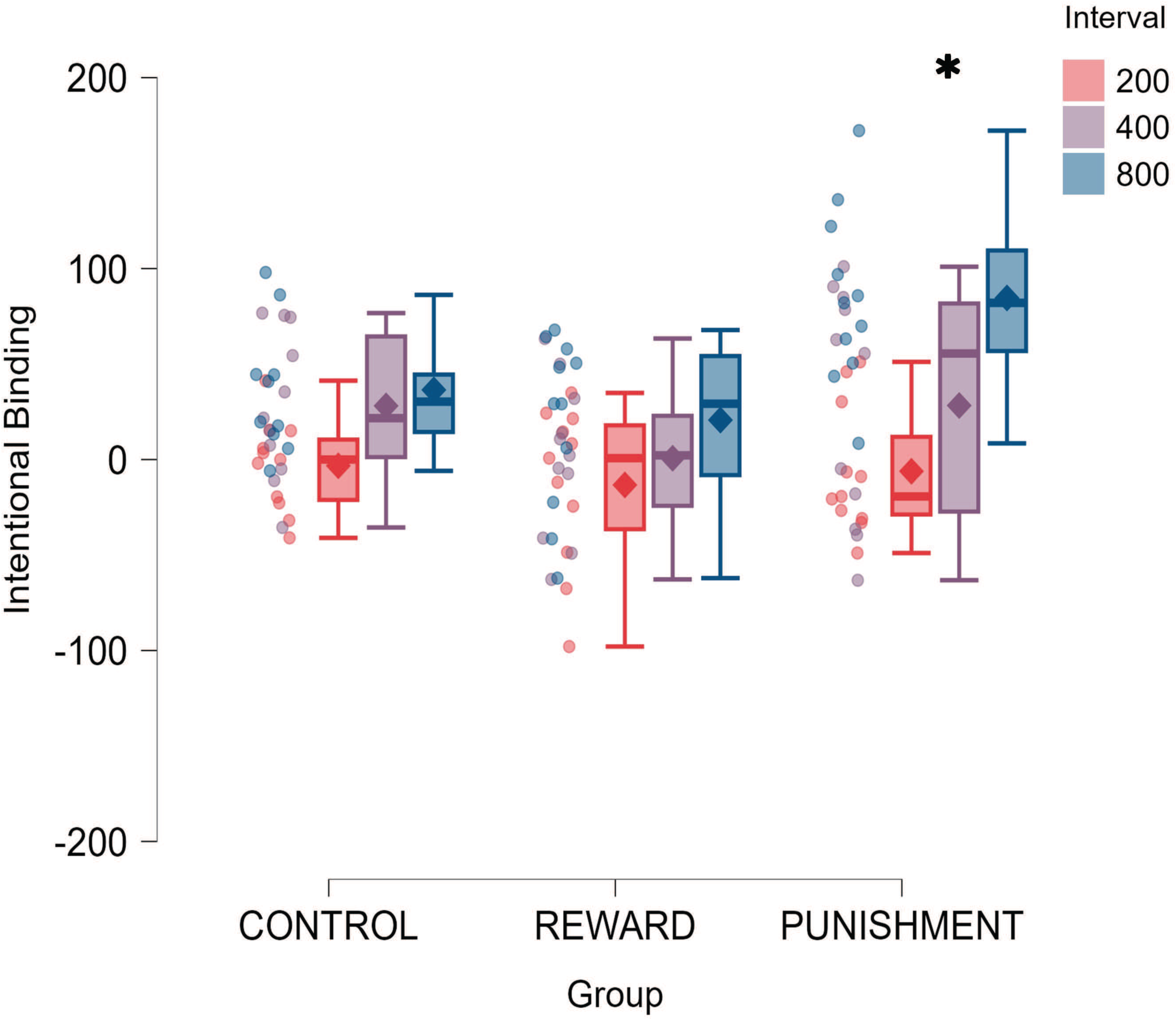
Boxplots for Reward, Punishment, and Control groups’ Intentional Binding for across the three time intervals [200ms (pink), 400ms (purple), 800ms (blue)]. Diamonds represent means and dots represent individual data. * = significantly different from Reward and Control.

### Outcome event-related potentials

Outcome ERPs for the Reward, Punishment, and Control groups in Active and Passive conditions in Fz, Cz, and Pz electrodes can be found in Fig 4.

**Fig 4:**
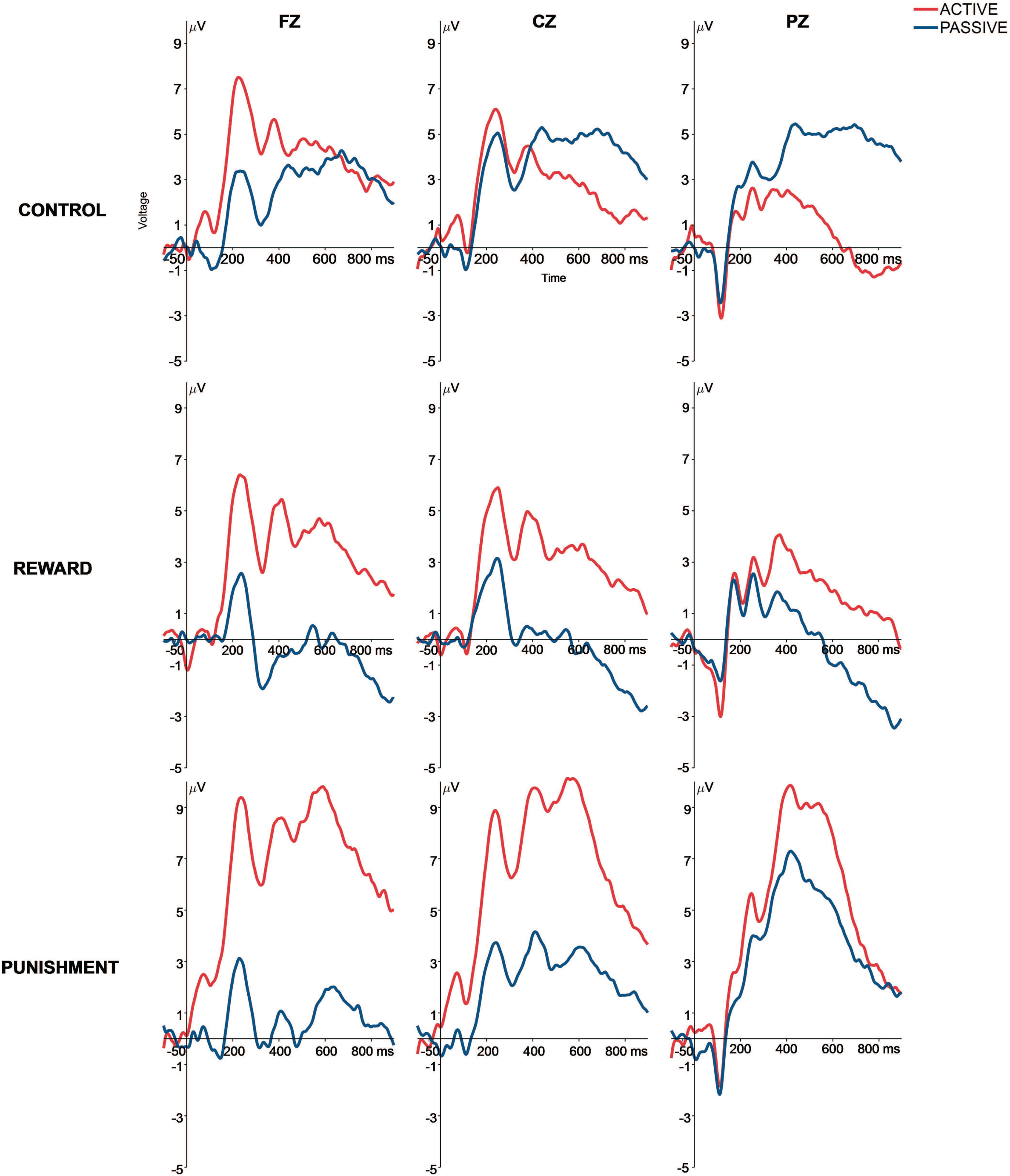
Outcome event-related potentials after judgement feedback presentation for the Reward, Punishment, and Control groups in the Active (red) and Passive (blue) conditions in the Fz, Cz, and Pz electrodes. Presented with a 30Hz low-pass filter.

#### N100 amplitude and latency

N100 changed between Active and Passive conditions in the Fz electrode (Fig 5). A significant main effect for Condition was found for N100 amplitude (F(2,84.192)=7.4316, p<0.001). The Active condition demonstrated a more positive N100 amplitude compared to the Passive condition [MD: 1.771, p<0.001, 95% CIs= 0.851-2.681].

**Figure 5:**
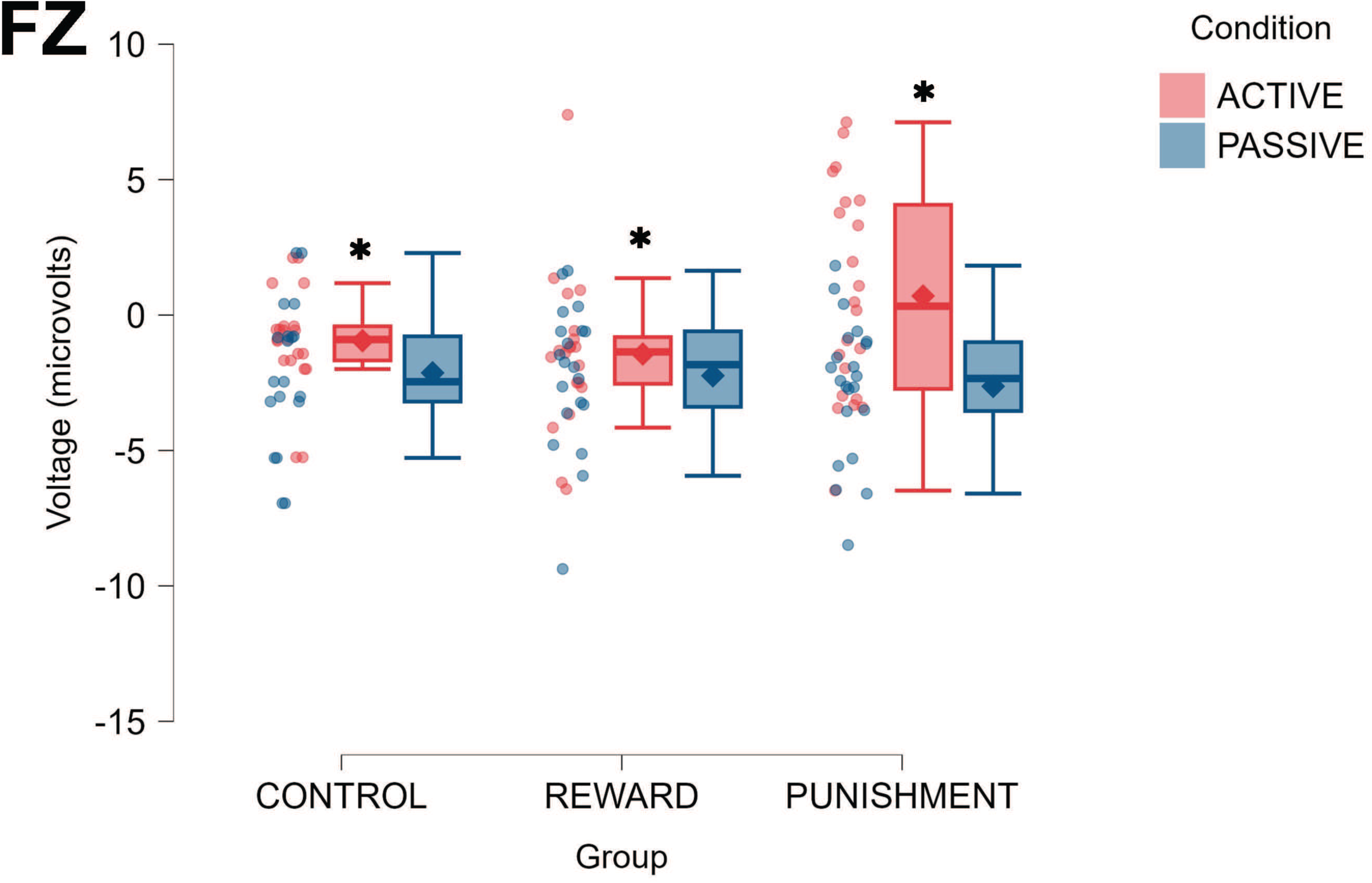
Boxplots for N100 amplitude in microvolts for the Reward, Punishment, and Control groups in the Active (red) and Passive (blue) conditions in the Fz electrode. Diamonds represent means and dots represent individual data. * = significantly different from Passive condition.

N100 latency was similar between groups and conditions (p>0.05).

#### P300 amplitude and latency

In the Pz electrode, Punishment demonstrated higher P300 amplitude regardless of the condition (Fig 6). A significant Group x Condition interaction was found for P300 amplitude (F(2,84.220)=4.067, p=0.021). Punishment elicited a higher P300 compared to Reward [MD: 5.554, p=0.04, 95% CIs= 0.192-10.915] and Control [MD: 7.131, p=0.006, 95% CIs= 1.781-12.480] feedbacks during the Active condition. Reward feedback elicited a higher P300 amplitude during the Active condition compared to Passive [MD: 2.421, p=0.038, 95% CIs= 0.140-4.702]. Punishment feedback also elicited a higher P300 amplitude during the Active condition compared to Passive [MD: 2.535, p=0.023, 95% CIs= 0.360-4.709]. No significant differences between Active and Passive conditions were found Control [MD: 1.161, p=0.192, 95% CIs= −3.836-0.783].

**Fig 6:**
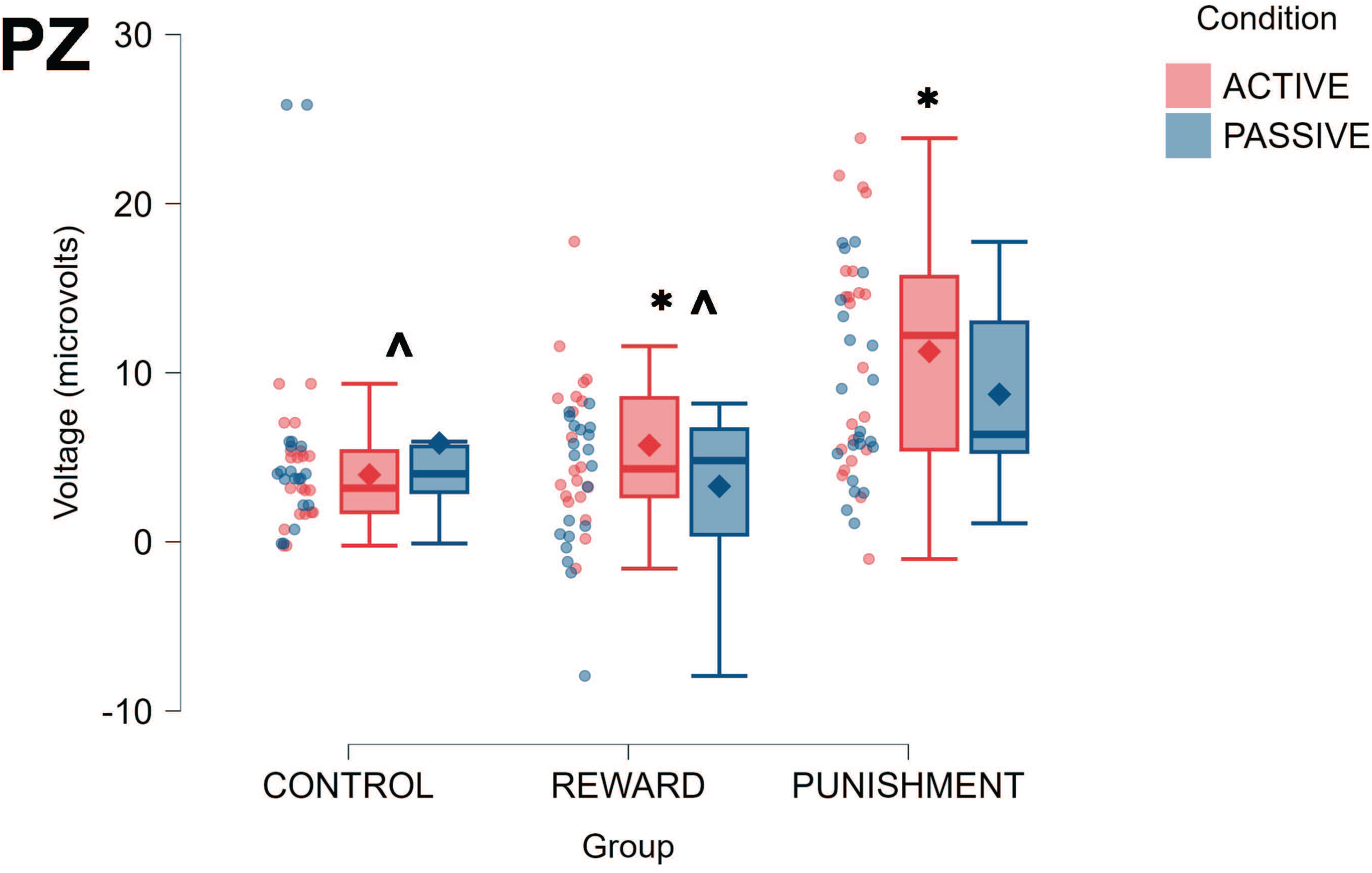
Boxplots for P300 amplitude in microvolts for the Reward, Punishment, and Control groups in the Active (red) and Passive (blue) conditions in the Pz electrode. Diamonds represent mean and dots represent individual data. * = significantly different from Passive. ^ = significantly different from Punishment.

Control feedback elicited a quicker P300 in comparison to reinforcement feedback. A significant main effect for Group was found for P300 latency (F(2,27.901)=5.077, p=0.013). Control demonstrated a faster P300 latency compared to Punishment [MD: 71.022, p=0.004, 95% CIs= 25.250-116.794] but not Reward [MD: 41.577, p=0.080, 95% CIs= 5.272-88.425]. No significant difference found between Reward and Punishment (p>0.05).

#### Late Positive Potential amplitude and latency

Punishment feedback changed brain activity more than Control and Reward feedbacks during the Active condition and compared to its Passive condition (Fig 7). A significant Group x Condition interaction was found for late positive potential amplitude (F(2,82.994)=5.076, p=0.008). Post-hoc pairwise comparisons revealed the Punishment group had a higher late positive potential amplitude during the Active conditions compared to Control [MD: 6.955, p=0.010, 95% CIs= 1.397-12.494] and Reward [MD: 6.243, p=0.024, 95% CIs= 0.680-11.805]. No differences were found between groups during the Passive condition (p>0.05). Punishment feedback also elicited a higher late positive potential amplitude during the Active condition compared to Passive [MD: 2.578, p=0.034, 95% CIs= 0.206-4.951]. No significant differences were found for Control and Reward (p>0.05)

**Fig 7:**
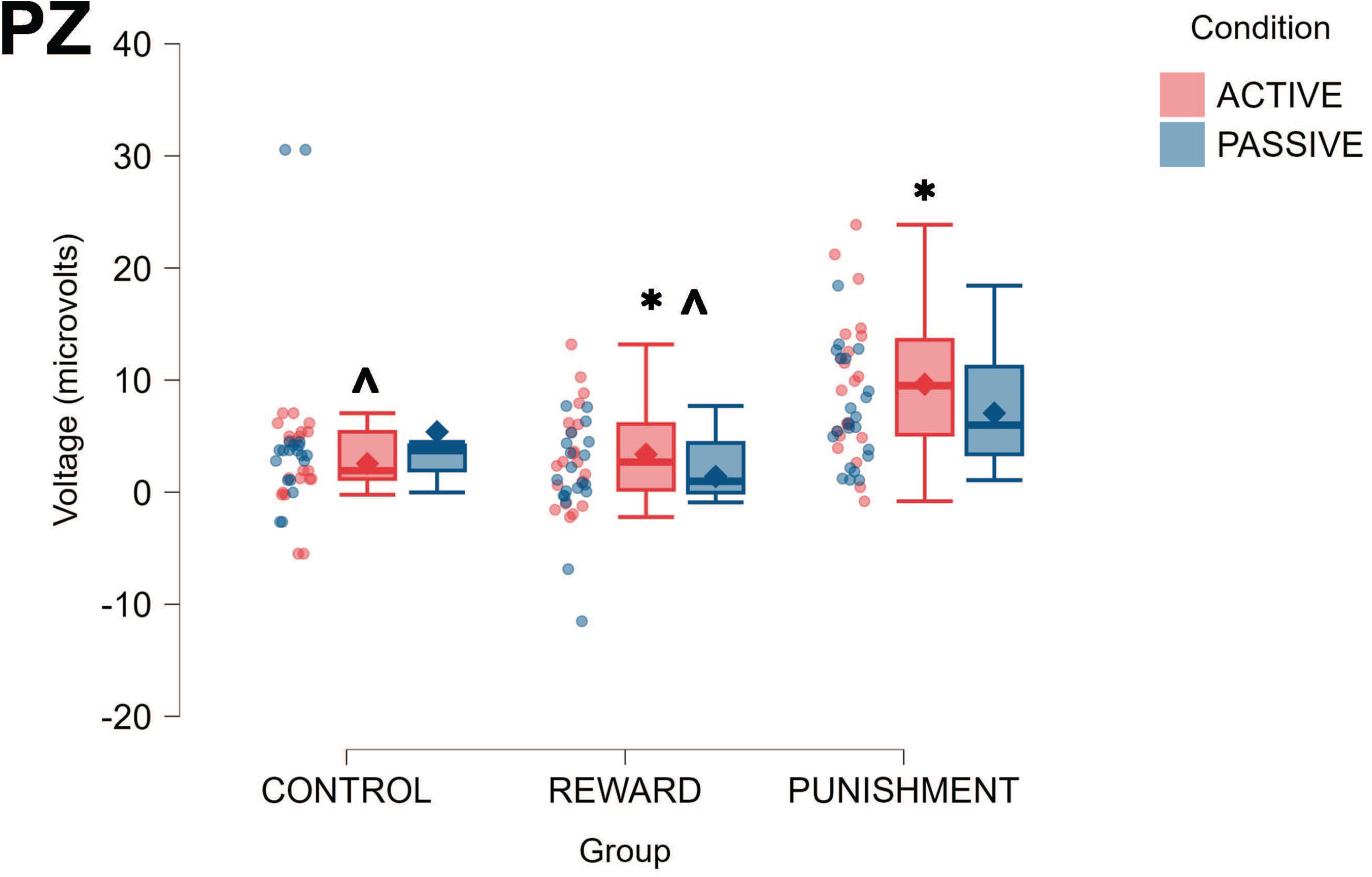
Boxplots for Late Positive Potential amplitude in microvolts for the Reward, Punishment, and Control groups in the Active (red) and Passive (blue) conditions in the Pz electrode. Diamonds represent means and dots represent individual data. * = significantly different from Passive. ^ = significantly different from Punishment.

Control feedback elicited a quicker late positive potential in comparison to reinforcement feedback. A significant Group x Condition interaction was found for late positive potential latency (F(2,84.602)=6.159, p=0.003). Control feedback elicited a faster late positive potential in the Active condition compared to Punishment [MD: 60.166, p=0.015, 95% CIs= 9.442-110.890] and Reward [MD: 68.220, p=0.006, 95% CIs= 16.288-120.152]. No significant difference found between Reward and Punishment (p>0.05). Passive condition elicited a slower late positive potential latency compared to the Active condition [MD: 75.690, p<0.001, 95% CIs= 41.690-109.690]

## Discussion

The purpose of this study was to evaluate how intentional binding was changed by modulating the valence of reinforcement feedback associated with temporal judgement accuracy and to examine the neural correlates underlying this feedback processing. We found that punishment feedback elicited greater intentional binding than reward feedback, but not more than control feedback. Punishment elicited greater ERP amplitudes in comparison to reward and control feedback, especially in the Active conditions. However, the punishment group’s ERP amplitude decreased from the Active to Passive conditions showcasing the role of agency in processing aversive feedback.

### Interval length and action type changed temporal judgement error

We found that judgement error was higher in longer intervals (∼800ms) than shorter ones (∼200ms,∼400ms). Moreover, judgement error was higher in the Passive condition compared to Active. These results are in line with other previous studies in this domain [66–70]. Longer intervals and actions performed by others limit the integration of embodied sensory cues (i.e. proprioception) to be entangled with external consequences. As a result, the brain’s forward models that generate a sense of agency are not adequately formed, leading to more error during longer intervals [69].

However, we did not find modulation of judgement error due to the type of feedback received. All groups had a similar amount of judgment error across the intervals. Previous studies in the context of skill learning, have found error is corrected more quickly when punished or rewarded [71–73]. This lack of an effect could be attributed to the randomized presentation of the intervals. By randomizing the tone intervals, the task became more challenging, thus harder to correct errors from one trial to the next.

### Punishment increased intentional binding

Previous studies have investigated outcome valence as a potential moderator of international binding. Aversive auditory tones associated with monetary loss decreased international binding compared to positive and neutral tones [10,11,74]. However, another study found no modulation of temporal binding with appetitive and aversive faces [75]. Di Costa et al., (2018) evaluated the effect of feedback valence after outcome judgements were made [12]. Importantly, this feedback was the same valence as the tone presented after their action and not in response to their judgement performance. They found greater international binding with negative feedback than reward feedback [12]. Our results align with this finding in that punishment feedback after the participants made interval estimations, increased intentional binding compared to reward, but was not significantly different from control feedback. We suggest that providing punishment via monetary loss, increases the binding of the action and outcome via greater attention and arousal [39,76]. Punishment feedback elicits more top-down control to avoid future punishments [39,76,77], in turn leading to greater action and outcome binding.

Intentional binding was not different between control and reward feedback, which aligns with Yoshie & Haggard, (2013). This indicates reward and control feedbacks create similar binding, despite the different feedback valence. It should be noted that our feedback presentation was different compared to these previous studies [10,11,74]. Instead of modulating the valence of the tone, we modulated feedback associated with the participant’s judgements (i.e. estimating the time between the button press and tone). This distinction in methodology could explain our divergent results and requires further testing.

### Punishment after self-initiated actions attenuated early outcome event related potentials

N100, in part, represents immediate error and sensory processing [42,60]. Previous studies have evaluated the effect of self- vs externally generated actions on outcome processing, using N100 [7,8,78–81]. These studies show N100 amplitude decreases during self-generated outcomes, reflecting sensory attenuation. Other studies have compared the effect of different outcome valences on N100 amplitude when comparing self and externally generated actions. One study observed no effect of outcome valence (positive or negative) on N100 amplitude during self-generated outcomes [82]. Our findings agree in part with these previous studies. Punishment feedback elicited a greater, N100 amplitude during the Passive condition compared to the Active condition, suggesting a sensory attenuation in error processing when pushing the button themselves. The amplitude attenuation during the Active condition can be attributed to the predictability of the outcome. A previous study noted N100 amplitude was not modulated by level of agency when outcomes were equally predictable [83].

During the Passive condition, participants were unable to form an adequate model of the task, as they did not actively press the button generating the tone. As a result, they are missing out on critical sensory information to form a complete model of the task. When feedback is then delivered in the Passive condition, the outcome is not as predictable, leading to a larger N100. Moreover, this effect is heightened with punishment feedback creating, a greater N100 when feedback is less predictable. This is similar to another study where N100 amplitude increased when punishment was delivered by a third-party [84]. However, it should be noted that other studies have shown decreases in N100 amplitude when actions were caused under coercion (i.e. low agency) [30] and no differences between active and passive actions [26,27]. Thus, more research is needed to understand how N100 is modulated by self- and externally generated actions.

### Reinforcement feedback increased late outcome event related potentials amplitude and latency

P300 represents various aspects of human perception, including attention, significance, and valence [51,58,85]. In the context of agency, P300 increases with high agency compared to low agency conditions [31,86–91]. Our findings align with this assertion, but as a function of feedback type. Reward and punishment feedback modulated P300 amplitude more in the Active condition compared to Passive and elicited a greater P300 compared to control feedback. Other studies have noted similar outcomes where P300 increased in response to a reward presented during an active task compared to a passive one [88,89]. Previous work suggests P300 reflects goal directed behavior and activity of the locus coeruleus [58,92–94]. We suggest reinforcement feedback activates the motivational circuits of locus coeruleus more during the Active condition than the Passive condition. However, it should be noted that, Majchrowicz et al., (2020) found no modulation of P300 with positive or negative outcomes during an interval estimation task [95]. This study evaluated the feedback valence associated with the outcome rather than feedback related to the participants interval judgement, as in this study.

P300 peak latency differed between groups. Control feedback elicited faster P300s compared to reinforcement feedback. Faster P300 latencies represent quicker stimuli classification [51,52] and lower cognitive load [96,97]. Reinforcement feedback includes information regarding valence and direction of performance, which inherently increases the processing time and cognitive load placed on the brain. This amount of information and context is not present in control feedback, resulting in slower processing of reinforcement feedback.

The late positive potential has been linked to affective processing and the activation extent of appetitive or aversive motivational systems [58,94,98]. Previous studies found increased late positive potential amplitude when viewing unpleasant images compared to pleasant or neutral [56,59,98,99]. Moreover, the late positive potential in response to aversive and appetitive images have divergent neural substrates. For instance, aversive stimuli recruit ventrolateral prefrontal, insula and cingulate cortex. Whereas appetitive stimuli recruit prefrontal cortex, amygdala and precuneus [100]. Punishment feedback increased the amplitude of the late positive potential in comparison to reward and control feedbacks, which can be indicative of punishment feedback engaging different brain pathways, thus increasing late positive potential amplitude. Moreover, these effects were attenuated during the Passive condition, with punishment feedback demonstrating similar late positive potential amplitude as control feedback. Stimulus significance is a driver of late positive potential amplitude [56,59,98,99]. Our findings signify that punishment feedback when delivered after actions not executed by the person themselves, loses significance and value compared to when they actively engage in the task. This mirrors our behavioral findings; a greater intentional binding was found for the punishment group (i.e. less judgment error in the active condition compared to passive).

Like our P300 results, latencies for peak late positive potentials were faster for the control feedback compared to reward and punishment groups. In part, late positive potential represents emotional processing of stimuli. Given that control feedback has non-valanced information, it does not recruit areas of the brain associated with emotional processing. Thus, resulting in quicker processing compared to reinforcement.

## Conclusion

In conclusion, reinforcement feedback based on temporal judgement accuracy changes our sense of agency and cortical processes associated with outcome monitoring. Specifically, we found that when we perform actions ourselves, feedback associated with our judgements are perceived as being more salient and significant, opposed to actions generated by others. Moreover, aversive feedback (i.e. punishment) enhances this effect compared to more appetitive or neutral feedback. These findings could be important for understanding changes to human sense of control and agency in psychiatric disorders, such as schizophrenia and obsessive-compulsive disorder.

